# Axonal distribution of Layers IV-VI Connections in Human Visual Cortex Revealed by in-vitro extracellular injections of Biocytin

**DOI:** 10.1101/2025.03.18.643885

**Authors:** Gad Kenan, Rafael Malach

## Abstract

The organization of interlaminar axonal connections was studied in slices of adult human visual association cortex. Cortical tissue was obtained during surgical glioma treatment in three subjects. Slices were injected with the highly sensitive anterograde tracer biocytin. In some cases, tract tracing was combined with laminar recordings of electrically evoked potentials, employing current-source density analysis. This study is based on 23 injections placed at various cortical layers and analyzed in detail. The results reveal a striking columnar architecture of local axons organized in vertically oriented bundles as well as vertical columns of neuronal cells bodies. A high degree of specificity was observed in the connectional patterns of various layers. This specificity was reflected in the tendency of lateral connections to spread into different cortical layers, and in the large variations in the axonal bouton densities in different layers. More specifically. Layer IV injections resulted in a more circular projection pattern including massive projections into layer II-III. Layer V injections produced extensive intra-laminar lateral connections while layer VI injections resulted in a vertical projection leading up to layer IV and a lateral intra-laminar projection. Thus, our results reveal highly specific connectional patterns in the different cortical layers in adult human association visual cortex.

## INTRODUCTION

The understanding of the human cortex provides a fundamental challenge in neurobiological research. Clearly, the attempt to relate cognitive, perceptual and motor functions to brain mechanisms will ultimately depend on a better knowledge of the human cortical circuitry. Unfortunately, due to technical difficulties involved in studying post-mortem material, the study of neuronal connections in the human cortex has been limited.

Most of the information on human cortical connectivity gained so far from the Golgi impregnation technique (Cajal 1899; Golgi 1873, Lorente de No, 1934) which suffers from short range axonal impregnation, and can be applied only to newborn and infant material. It is of course a standing question how valid the extrapolation from infant material is to fully developed adult tissue. For example, in monkey, synaptic connection of upper cortical layers is rapidly eliminated and modified up to puberty (Borgeois and Rakic, 1993).

More recently, studies have examined human cortical tissue slices obtained following clinical surgical procedure to study local synaptic connectivity between human cortical neurons (Luke Campagnola et al 2022, Hunt et al 2023, Thomson and Bannister 2003, Bannister 2005). Here We describe the extracellular application of highly sensitive tract tracing method, using the tracer biocytin (Horikawa and Armstrong, 1988) in the in-vitro human slice preparation (Kenan-Vaknin et al., 1992a; Kenan-Vaknin et al., 1992b). The tracer biocytin is rapidly transported (Malach et al., 1993) and combines high sensitivity with Golgi-like labeling of axons and neuronal somata (Yoshioka et al., 1992; Kenan-Vaknin et al., 1992a; Malach et al 1993, Amir et al., 1993). The range over which the axons can be traced is currently limited only by the size and thickness of the slices and may be extended by modifications in slice incubation parameters. Mohan et al (2015), employing intracellular biocytin staining of dendrites and axons in human cortex described the axonal distribution of 3 cells in layer IV extend to layers II-III and V-VI. The axonal distribution of cells in layer V, extend to layer V and IV. Campagnola et al, (2022), generated a comprehensive dataset describing synaptic connections within each layer in the mouse and human cortex.

In the present report we aimed at elucidating the interlaminar distribution of axonal connections in the adult human visual association cortex, Layers IV, V and VI. Studies in primates have a revealed a powerful interlaminar connectional route (Cajal, 1899; Lorente de No, 1949; Lund and Booth, 1975; Rockland and Lund, 1983; Livingstone and Hubel, 1984; Yoshioka et al., 1992, Thomson and Bannister 2003, Bannister 2005)

An important feature of these connections is their laminar specificity, with individual cortical layers having pattern of connections which are unique in their target layers, the extent of arborization and the lateral spread within individual layers. Considering only the major axonal interlaminar pathways, these studies suggest a basic pattern of interlaminar connectivity whose main course of origin lies in layer IV from which axons project to upper cortical layers. From supragranular layers the axonal projections reciprocally connect mostly to and from infragranular layers.

Our results are compatible with this picture and extends it to reveal an intricated and extensive set of specific functional interlaminar connections in visual association cortex of the adult human. In the present manuscript we focus on layer IV-VI connectivity. The connections of upper layers were described elsewhere Kenan-Vaknin et al., 1992b.

## METHODS

### Human cortex slice preparation

Human cortical tissue was removed during neurosurgical treatment for gliomas in three adult subjects (1-5cm^3^) were removed suction (one subject) or a cut using a scalpel (two subjects). In all cases the tissue was taken from the posterior portion of the dorsal bank of the occipital lobe. Based on Nissl preparations and the gyral pattern at the site of surgery, the area under study was most likely the peristriate region of the visual cortex. In most cases, the tissue used for analysis appeared to be normal (i.e. no abnormal infiltration of blood vessels, and as indicated by Nissl staining). In a few slices, abnormal blood vessels were seen at the edge of the slice but did not affect the staining. Following surgery, the tissue was immediately placed in oxygenated, cold (0-5°C) artificial cerebrospinal fluid (ACSF, containing in mM: NaCl 124, KCl 4, NaH_2_PO_4_ 1.25, CaCl_2_ 2.5, MgSO_4_ 1.5, NaHCO_3_ 26, at pH 7.4). The tissue was transferred to a laboratory next to the operation room and coronal and horizontal slices (400um thick) were cut using a McIlwain tissue chopper and placed in three interface recording chambers which were modified (large pool, 5cm in diameter at 34°C for 2h prior to staining and recording.

### Tract tracing

We used 2.5% biocytin (Molecular probes, Eugene, Oregon) dissolved in 0.05M Tris buffer pH 7.6, applied iontophoretically (3-10uA for 3-10min) using micropipettes (10-15um in diameter) mounted on a digital micromanipulator. Micropipettes were guided under direct observation through a dissecting microscope, 100-200um deep into the slice. One injection was made in each slice. Thirty-eight injections were made, and 23 focal and well stained injections were used for this study. Injections were mad in layers II-III (n=5), IV (n=5), V (n=3), VI (n=4) and 5 injections in the middle of horizontally cut slices.

### Recording of field potentials

Extracellular laminar field potentials were evoked by distant layer IV stimulation (400um away from the recording track, 10-15v square pulse), using concentric bipolar stimulating electrode, minimizing electrical spread of current (Chiaia and Teyler, 1983). In many cases a dark line of fibers corresponding to layer IV was seen under transillumination of the slices, enabling a precise localization of the stimulating and biocytin micropipettes. Extracellular recordings were made every 100um along the radial axis (from pia to white matter). Recordings were obtained with microelectrodes pulled from 1.2mm glass pipettes (tip diameter < 1um) filled with 3M NaCl. One dimensional Current-source density analysis was employed (Nicholson and freeman, 1975) using the extrapolation method for complete description of the laminar distribution of current densities (Vaknin et al., 1988). Electrophysiological recordings were combined with biocytin injections in layer IV.

### Histology

After injection, the tissue was left for 1-8hrs. and then fixed with 4% paraformaldehyde, overnight. Sixty um sections were cut from each slice and were processed according to Horikawa and Armstrong (1988) with some modifications (Kenan-Vaknin et al., 199a). One section from each slice was stained for Nissl. In some cases, slices were put in fixative immediately after cutting and then stained for Nissl. Sections were inspected at high and low power using bright- and darkfield microscopy.

## RESULTS

### Axonal connections form bundles

A striking observation that appeared to be invariant to the laminar location of the injections is the consistent grouping of axonal connections along the vertical dimension. Prominent vertical bundles of axons were a dominant feature and are illustrated both schematically and in micro-photographs in figure 1.

**Fig. 1.**
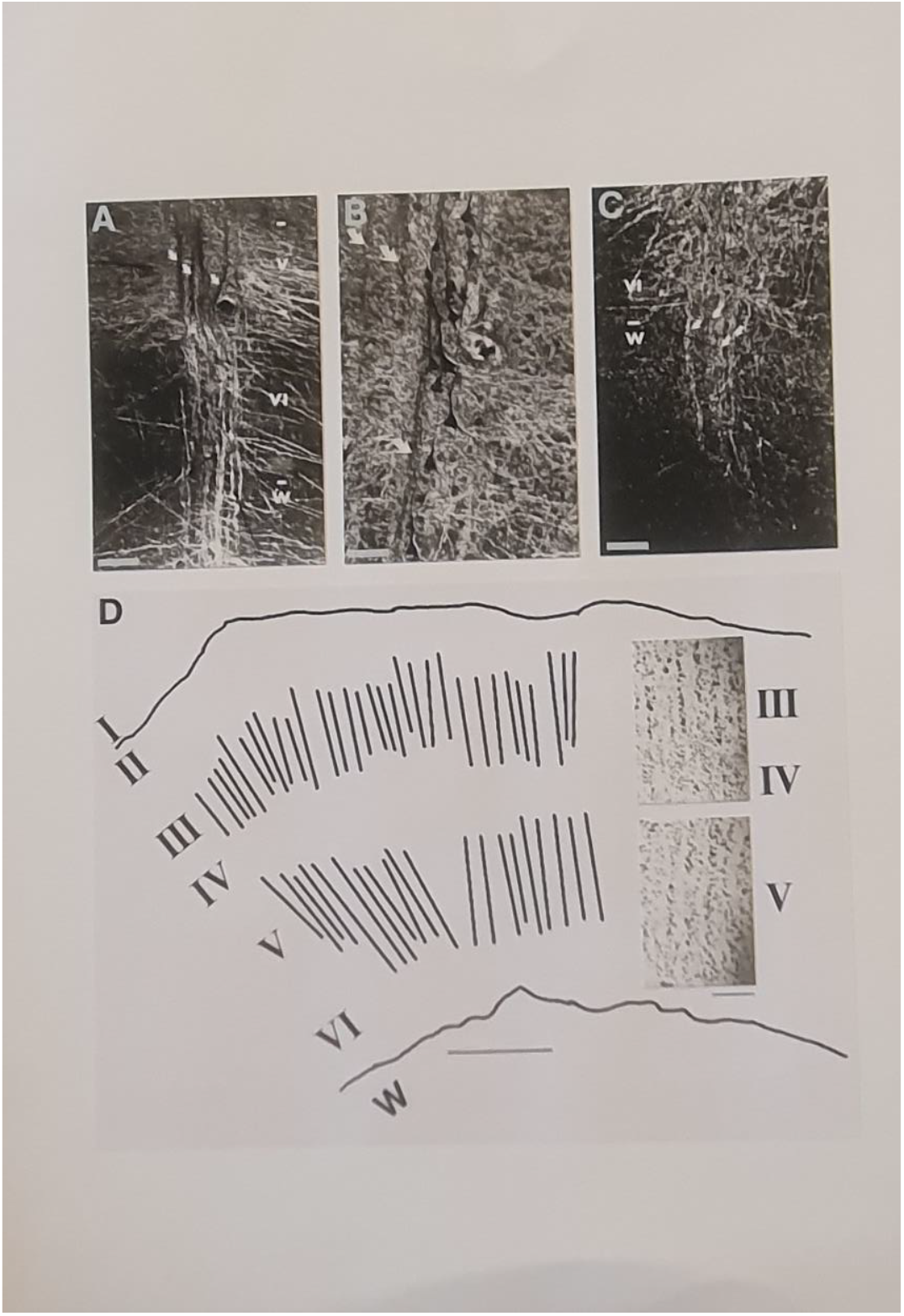
columnar organization of bundle of axons (a) and cells (b) as depicted in both biocytin and Nissl stain (d)

#### Axonal connections of layer IV

Injections in layer IV were placed under visual guidance by targeting the dark line of fibers in this layer which could be visualized by transilluminating the slice. Figure 2 is an example of such injection. Note that the axonal projection pattern here is in a striking contrast to that produced by layer II-III injection. The wide lateral spread of axons found in layer II-III injections is lacking, instead a more circular axonal projection pattern is evident, with massive projections into layers II-III and fairly short fibers running in layer IV. Few fibers could be observed extending further away in layer IV and vertically up to layers I-II, and few to layer VI.

**fig. 2.**
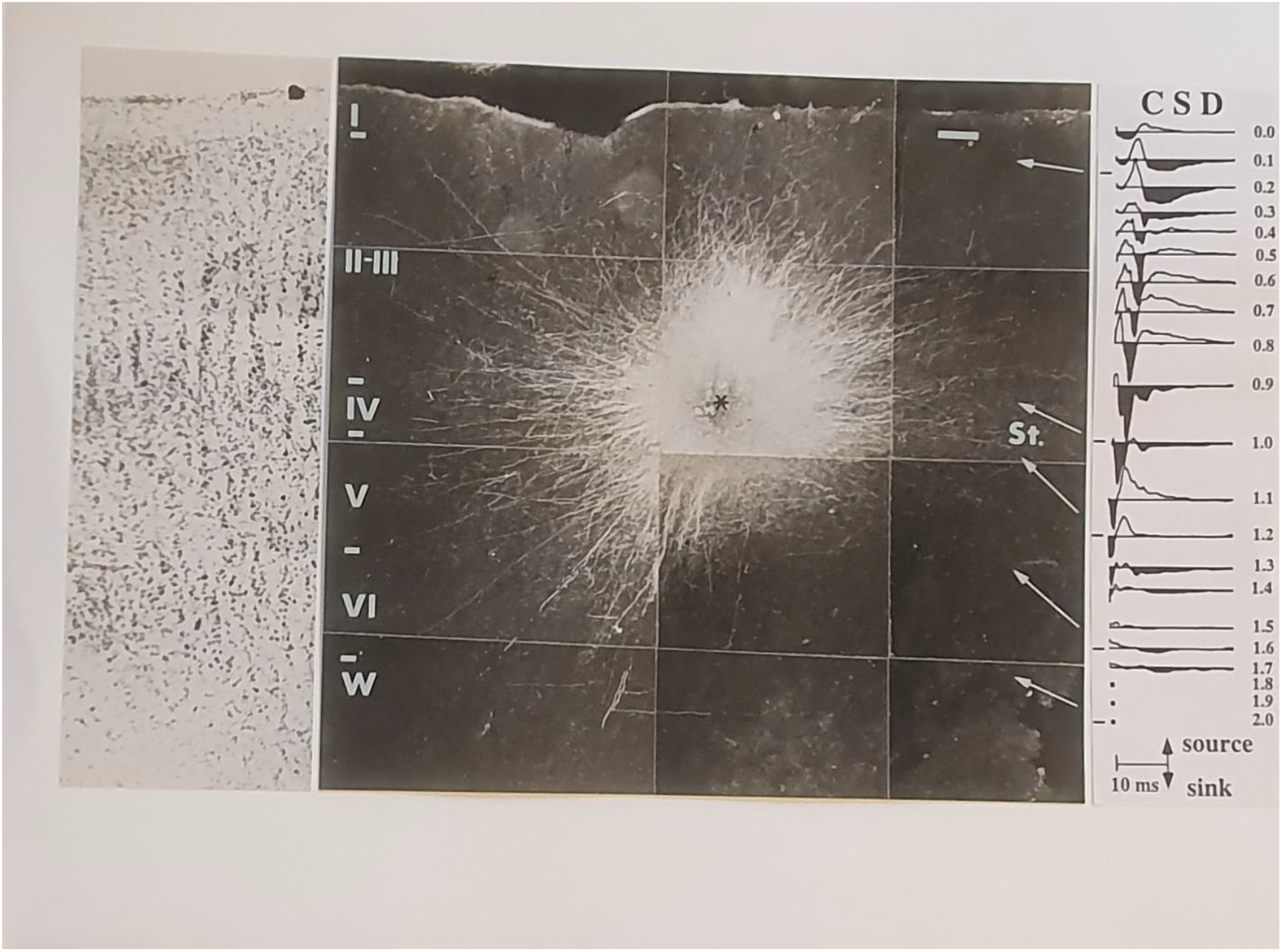
layer IV injection, a circular axonal projection pattern is evident, with massive projections into layers II-III and fairly short fibers running in layer IV. Few fibers could be observed extending further away in layer IV and vertically up to layers I-II, and few to layer VI. CSD analysis of layer IV stimulation, reveals activation of the entire cortex with most of the activity taking place in the supragranular layers.

#### Electrophysiological study of layer IV projections

To assess the functional impact of layer IV projections, electrical stimulation was delivered to layer IV under direct visual guidance. Multiple recordings of the local field potential were performed along the radial axis 400um away from the stimulating site. Following recordings, current-source density analysis was applied to reveal the spatio-temporal distribution of synaptic currents. The results (fig. 2, CSD, right panel) show that electrical stimulation in layer IV leads to early, short duration and high amplitude sinks in layers I-V, followed by lower amplitude, longer duration, sources in layer II-III. The earliest sinks appear in layers I, IV and V. starting in layer V, latencies of sinks increase at more superficial layers. The corresponding sources are located in layers I-III and IV-V. Late activity is seen in lower layer III-VI. In summary, layer IV stimulation resulted in activation of the entire cortex with most of the activity taking place in the supragranular layers.

#### Axonal connections of layer V

Injections in layer V produced yet another axonal pattern which is shown in fig. 3. Here, similar to the layer II-III injections, axons could be found distributed extensively in the horizontal dimension. Unlike injections in supragranular layers, Layer V axons are mostly confined to layer V. Many oblique running fibers could also be observed, coursing either upward towards layer I and down to layer VI. Along the vertical axis, bundles of axons could be seen coursing towards the white matter (fig. 1c) and upwards towards supragranular layers.

**Fig. 3.**
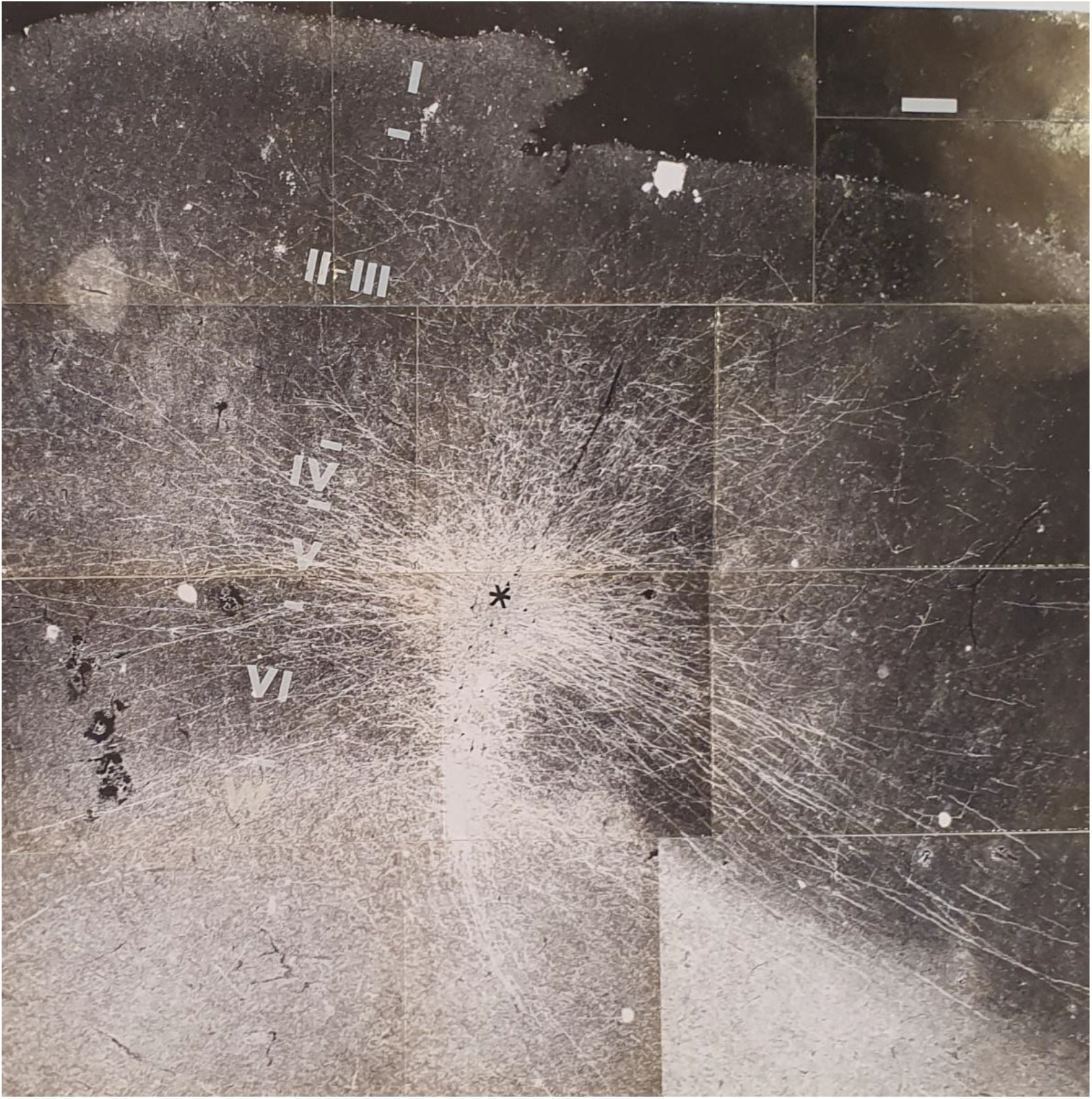
Layer V injections produced extensive intra-laminar lateral connections and fibers exiting to whit matter.

#### Axonal connections of layer VI

Figure 4 shows an example of an injection placed within layer VI.

**Fig. 4.**
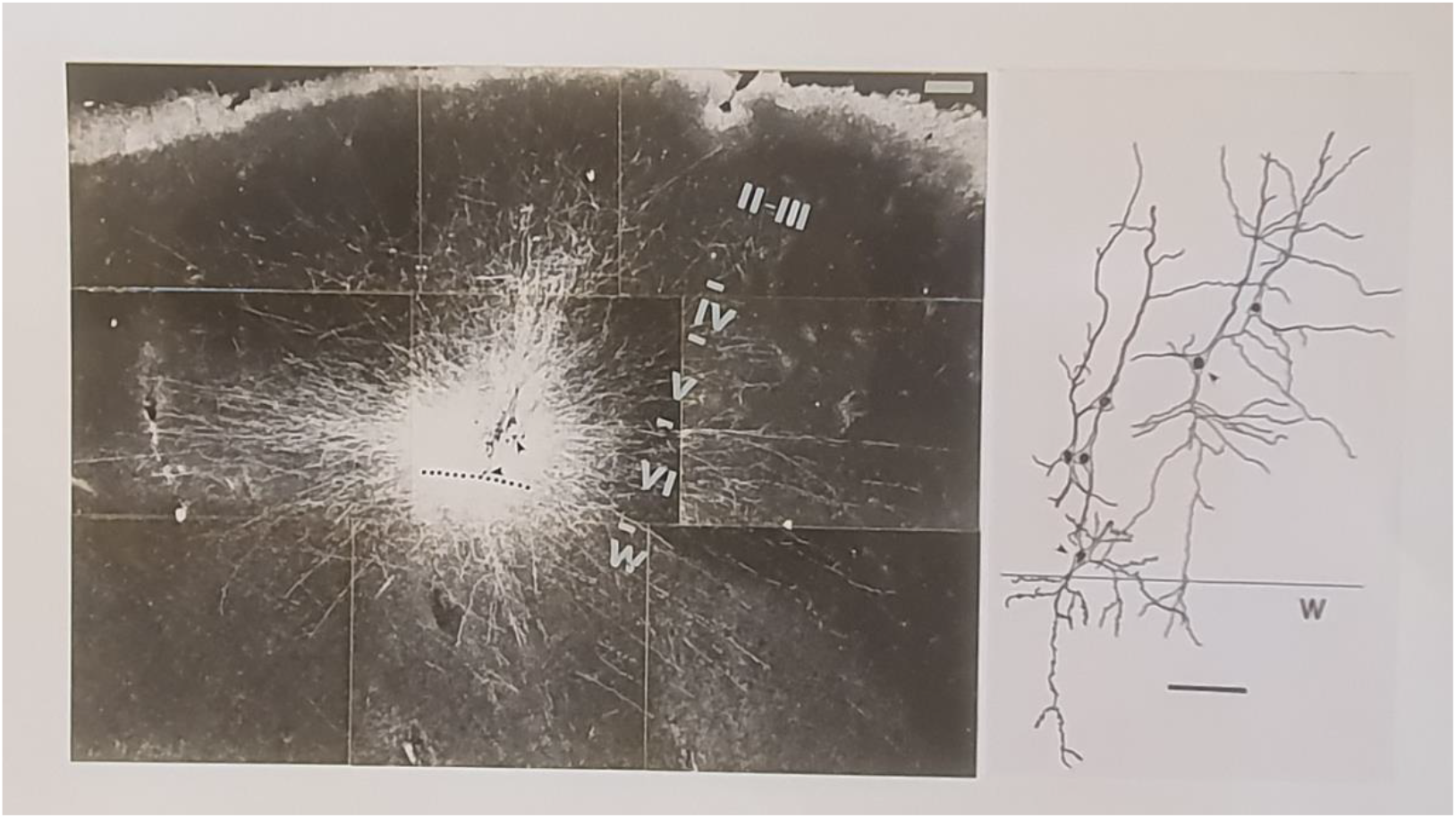
layer 6 injection, fibers mostly confined to layer 6 in the lateral dimension.

Extensive and focused vertical projections could be observed reaching up to layer IV. Additional, weaker projections appeared to continue upward towards layer I. In the lateral direction, fibers were confined mostly to layer VI. Obliquely running axons appeared to radiate in all directions and ascended up to layer I and down to white matter. Quite often we could observe “inverted pyramids” in this layer with descending dendritic shaft directed towards the white matter. Dendrites of neurons situated close to the white matter ignored the white /gray matter boundary and extended spine bearing dendrites into the white matter (fig. 4, right panel)

#### Axonal structures

We have observed numerous cases in which axons radiating away from the injection site suddenly changed course or branched more extensively at some cortical points. Also, the hallo around the injection site was often organized in tongue like formations suggesting that some cortical regions around the injection site were preferred targets for the axonal projections. However, due to size limitations of the slices examined-we were not able to trace the axons at sufficient distance to reveal the lateral patches or cluster of connections found in monkey cortex (e.g. Amir et al., 1993)

## DISCUSSION

### Methodological problems

Although the tracing method described here provides highly detailed connectional information, it does nonetheless, have two limitations which should be considered. One problem inherent in the method is the likelihood of “false negative” (i.e. failure of labeling of existing axons). This problem stems from two possible reasons: first, in preparing the slices, axonal projections which do not run precisely within the section will be severed. The likelihood that axons will deviate from the section and be cut increase substantially with the distance from the injection site. Second, for effective transport to occur, the neuron has to be kept viable for 3-6 hours. Clearly, many neurons can be maintained for such time periods as evinced by the axonal labeling and physiological activity, but it is likely some neurons are lost. We do not have a reliable estimate of what proportion of neurons remained viable throughout the procedure. Both problems results in an underestimate of the connectional spread and this should be born in mind when interpreting the data.

Another problem is the lack of clear delineation of cortical areas' boundaries in our set-up. Consequently, injections that may have been placed in different cortical areas were lamped together in this study. Nevertheless, it appeared that the overall pattern of connections remained quite invariant across injections. It remains to be seen if more detailed, quantitative analysis of parameter such as spread of connections, the size of axonal halo etc., may prove to vary systematically in different injected areas as they do in macaque (Amir et al 1993).

### Laminar specificity

A prominent feature of the axonal connections was their tendency to show differential targeting and spread across laminae. This phenomenon was clear cut in the lateral spread of axons, so that in certain laminae the axonal projections spread over long distances, while at others they remained confined. This was particularly striking in the axonal projections following layer II-III injections (Kenan-Vaknin et al 1992), where axonal projections clearly avoided layer IV and VI, and arborized extensively in layers II-III and V Similar, laminar specific spread was observed following layer V and Layer VI injections (fig. 3 and 4 respectively) except that here the majority of lateral connections remained intra-laminar.

When viewing the vertical axonal distribution from this perspective the sequence of the main vertical projection appears to be as follows: from layer IV upward to the bottom half of layers II-III. From layer II-III down to layer V. From layer V-VI the vertical ascending projections are quite diffuse but seems most prominent to layers IV and II-III.

It should be noted that we did not find evidence for subdivisions within the individual layers. However, the large size of injections used may have obscured specific labeling originating or targeting distinct sublayers.

In Summary, it is clear that there is laminar specificity both in the preferential lateral spread of axonal projections, in targeting specific lamina by vertical directed axons, and in the unique “signature” of the connectional patterns exhibited by axonal projections emanating from each layer.

